# Head model personalization: A framework for morphing lifespan brain images and brains with substantial anatomical changes

**DOI:** 10.1101/2021.04.10.439281

**Authors:** Xiaogai Li

## Abstract

Finite element (FE) head models have emerged as a powerful tool in many fields within neuroscience, especially for studying the biomechanics of traumatic brain injury (TBI). Personalized head models are needed to account for geometric variations among subjects for more reliable predictions. However, the generation of subject-specific head models with conforming hexahedral elements suitable for studying the biomechanics of TBIs remains a significant challenge, which has been a bottleneck hindering personalized simulations. This study presents a framework capable of generating lifespan brain models and pathological brains with substantial anatomical changes, morphed from a previously developed baseline model. The framework combines hierarchical multiple feature and multimodality imaging registrations with mesh grouping, which is shown to be efficient with a heterogeneous dataset of seven brains, including a newborn, 1-year-old (1Y), 2Y, 6Y, adult, 92Y, and a hydrocephalus brain. The personalized models of the seven subjects show competitive registration accuracy, demonstrating the potential of the framework for generating personalized models for almost any brains with substantial anatomical changes. The family of head injury models generated in this study opens vast opportunities for studying age-dependent and groupwise brain injury mechanisms. The framework is equally applicable for personalizing head models in other fields, e.g., in tDCS, TMS, TUS, as an efficient approach for generating subject-specific head models than from scratch.

## 1 Introduction

Finite element (FE) head models have become powerful tools in many fields of neuroscience, such as brain stimulations using direct current (tDCS) [1–8], magnetic (TMS) [9], and ultrasound (TUS) [10]. FE head models are also being used to study the development of neurodegenerative diseases [11–13], biomechanical consequences of neurosurgery [14–19]. In particular, FE head models have been tremendously used to study traumatic brain injuries (TBIs) in the last decades (see review [20–22]). TBI is still a leading cause of injury-related death and disability [23], causing a substantial threat to global public health and an enormous economic burden for society [24]. FE head models have played an important role in revealing the biomechanics and potential injury mechanisms of TBI and for developing prevention strategies.

Mesh generation is a prerequisite for FE analysis by discretizing a continuous domain into a finite number of elements, e.g., tetrahedral or hexahedral elements. Generation of FE head meshes is often time-consuming and challenging due to the complex geometry, though tetrahedral elements are relatively easier to generate, e.g., 1-2 hours [3, 4] for generating head models applicable for tDCS, TMS, and TUS. Such efficiency is greatly attributed to the well-developed automatic tetrahedral meshing algorithms within mathematics and computer science [25]; it is also due to the involved partial differential equations (PDEs) are less computationally demanding, which permit a huge number of tetrahedral elements (up to >10 million [1]) to capture brain’s complex geometry. Thus, personalized simulations with detailed and subject-specific head models are largely facilitated in brain stimulation fields.

Hexahedral elements are much more challenging to generate but are preferred for FE head models intended for studying TBIs (hereafter called head injury models) due to the significant advantages in accuracy or efficiency. The involved PDEs in FE head impacts consist of geometrical, material nonlinearity, and complex contacting algorithms, which is much more computationally demanding than brain stimulation fields. As a result, many head injury models [26–33] adopt conforming hexahedral elements with brain simplified. For example, these models have smoothed out brain surfaces without sulci, and gyri, resulting in fewer elements (often < 1 million). While sacrificing anatomical biofidelity in these models is partially a tradeoff for computational efficiency and is also due to the much less developed automatic computer algorithms for hexahedrons [25, 34]. Worth mentioning that voxel approach is efficient in generating (non-conforming) hexahedral elements by converting image voxels to hexahedral elements, either directly or with smoothing algorithms. However, a known concern is a less accurate peak strain/stress predicted, especially at the surfaces due to jaggedness, thus is not preferred. Most state-of-the-art head injury models use conforming hexahedral elements despite the meshing challenges [35]. Besides the less developed meshing algorithms for hexahedrons, a necessity to include falx and tentorium due to the important structural influence on brain mechanical responses under an impact [36] poses an additional challenge for subject-specific head injury model generation; while these structures are often neglected in head models for tDCS, TMS and TUS. A detailed analysis of the current meshing challenge for head injury models is found in a previous study [35].

Generation of FE head injury models with anatomical details remains a challenge and has become a bottleneck hindering personalized simulations. FE head models without anatomical details such as sulci and gyri also hinder studying detailed mechanisms at areas of interest, such as CTEs with pathologies observed at sulcal depth [37]. Studies have also shown the brain sizes/shape influences brain response significantly under impact [35, 38], suggesting the necessity to use personalized models to study the onset of TBI in real life. Along with the tremendous efforts being put on adult healthy FE head model generation, efforts on elderly, children, and infant models are less, being only a handful of models exists (e.g., [39–46]). The same or even higher challenge exists for generating these head models. TBIs are influencing all age groups, especially infants and elderly are over-represented [47], which urges the need for such models both to understand the injury mechanisms and to develop preventions. Thus, it is imperative to investigate efficient approaches for generating anatomically detailed subject-specific head models of lifespan suitable for studying TBIs.

This study addresses the challenge of generating personalized head models with conforming hexahedrons suitable for studying the biomechanics of TBIs, especially concerns about mesh morphing (also called warping). Mesh morphing is an approach for efficient generation of subject-specific models. It has been used extensively in many biomechanics fields on different organs [48–54], full-body models [55–57], as well as has for brain models with smoothed surfaces [18, 58-60]. A typical procedure includes image registration (rigid or affine and followed by nonlinear registration algorithms), from which a displacement field representing the geometrical difference between the subject and baseline mesh is obtained. Next, the displacement field is applied to morph the baseline mesh, resulting in a personalized mesh with updated nodal coordinates while remaining element connections. In general, the computed displacement field should comply with continuum mechanics conditions on motion, requiring diffeomorphic, non-folding, and one-to-one correspondence to avoid excessive element distortions [52].

In particular, deformable image registration-based mesh morphing has been shown promising to personalize healthy brain models due to its capacity to capture the anatomical difference between the baseline and the subject image [35, 61]. However, despite intensive efforts for decades, deformable image registration for subjects with substantial anatomical changes is still challenging and with limited registration performance [62]. When applying image registration for mesh morphing, an extra challenge is posed on the smoothness of the displacement field for morphed FE elements to remain acceptable element quality. Otherwise, without such reasonable element quality, not only an FE analysis is prevented from being carried out (e.g., when FE elements have negative Jacobian), also numerical accuracy is influenced. Therefore, although mesh morphing is efficient, one major challenge for using it to generate a detailed subject-specific FE head model is how to design an image registration pipeline that both leads to high registration accuracy meanwhile not causes excessive element distortions.

The aim of this study is to present a personalization framework capable of generating subject-specific head models within the whole lifespan and pathological brains with substantial anatomical changes (e.g., hydrocephalus brain, decompressive craniotomy, or TBI patients with squeezed ventricles). The framework consists of hierarchical multiple feature and multimodality registration steps, combined with mesh grouping. The feature steps are to handle substantial anatomical differences (e.g., large ventricle changes and brain lesions), allowing a good initialization before applying deformable image registration to further capture the inter-subject anatomical details; while the multimodality is to further improve anatomy alignment (e.g., T2W MRI images allows better alignment for CSF). To demonstrate the capacity of the framework, seven head models of lifespan, including a newborn, 1Y, 2Y, 6Y, adult, 92Y, and a hydrocephalus brain, are generated, showing its capacity for generating subject-specific models of all age groups, and pathological brains with large anatomical changes.

This study is organized as follows. In Methods, the images of the seven subjects are presented, followed by an overview of the framework, including the registration pipeline, mesh morphing, and mesh grouping. Matrices of Dice coefficient (DICE), Jaccard index, and 95th-percentile Hausdorff distance (HD95) are then calculated between the morphed image and subject’s T1W image to evaluate personalization quality. Finally, applications of the pipeline to the seven representative cases are detailed.

## 2 Materials and methods

### 2.1 Subjects

Seven subject-specific head models are generated to demonstrate the capacity of the proposed framework, including lifespan and hydrocephalus brain (**Fig. 1**). All images are acquired from previously published open-access datasets except the hydrocephalus brain is from the author’s previous study, and ICBM template is used as the template/fixed image for registrations and the baseline FE model creation. A summary of the brain images is provided in **Table 1**, briefly:

- Images of a newborn, 1Y, and 2Y are obtained from the UNC Infant 0-1-2 atlases [63] constructed based on 95 subjects with complete 0-1-2Y longitudinal scans of T1W and T2W images acquired with a 3T MRI scanner. The atlas has been reconstructed using infant longitudinal segmentation and groupwise registration techniques as detailed in [63]. Each atlas consists of T1W images, tissue probability maps, and anatomical parcellation maps.
- Image of a 6Y brain is obtained from the Pediatric atlases [64, 65]. The atlas was constructed using data from healthy children from the 4.5-8.5 age group using an iterative nonlinear co-registration algorithm as detailed in [65]. The atlas contains T1W, T2W, PD, and classified tissue volumes.
- Image of a single subject from the WU-Minn HCP database in the 26-30 age group, including both T1W and T2W image acquired with a 3T MRI scanner [66].
- Image of an elderly (92Y) from the Brain Imaging of Normal Subjects (BRAINS) atlas, which was created from 48 healthy elderly subjects within age group 91-93YO as detailed in [67]. The atlas contains T1W and tissue probability maps.
- Image of a hydrocephalus subject with a mass lesion at the brain stem front is reused from a previous study [68].
- The 1mm isotropic ICBM 2009c Nonlinear Symmetric template [64, 65] was constructed based on T1W images from 152 subjects between 18.5-43.5Y acquired on a 1.5 T MRI scanner. All images have been first linearly registered, and afterward, nonlinear deformations were iteratively estimated and generate the template as detailed in the original publication

**Table 1.** Subjects involved in this study.

| Subject | ICV (ml) | Imaging modality used | Image sources |
| --- | --- | --- | --- |
| 0-year-old | 463 | T1W (atlas) | [63] |
| 1-year-old | 1015 |  |  |
| 2-year-old | 1274 |  |  |
| 6-year-old<sup>1</sup> | 1842 | T1W (atlas) | [64, 65] |
| adult | 1480 | T1W, T2W (single subject in age group “26-30”) | [66] |
| 92-year-old<sup>2</sup> | 1323 | T1W (atlas) | [67] |
| Hydrocephalus | 1255 | T1W (single subject) | [68] |
| Baseline ICBM | 1885 | T1W, T2W (atlas) | [64, 65] |
| <sup>1</sup>Atlas of age group 4.5-8.5Y denoted as 6Y throughout this study for simplicity. |  |  |  |
| <sup>2</sup>Atlas of age group 91-93Y denoted as 92Ys throughout this study for simplicity. |  |  |  |

**Fig. 1.**
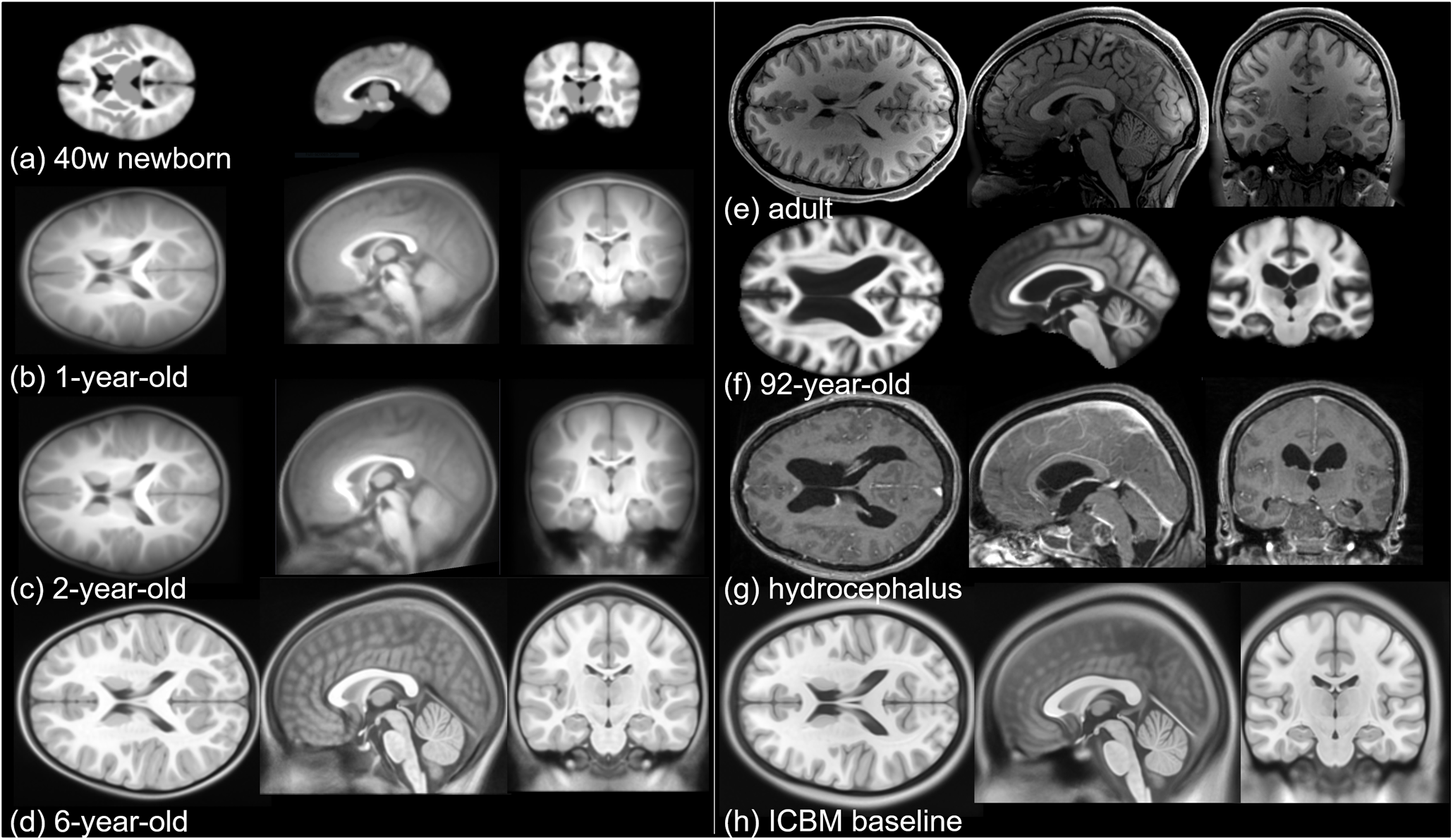
Image data used in this study. Axial, coronal, and sagittal views of **(a)** 40-week-old newborn, **(b)** 1-year-old, **(c)** 2-year-old, **(d)** 6-year-old, **(e)** an adult, **(f)** an elderly of 92-year-old, **(g)** T1W image of a hydrocephalus brain, and (**h**) the ICBM template corresponding to the baseline FE mesh to be morphed. For detailed preprocessing steps for these T1/T2 images, the readers are referred to the original studies.

### 2.2 Baseline FE head model

The baseline FE head model (the ADAPT model) [35] has been generated based on the ICBM template image and has the same geometry as the ICBM image. The ADAPT Fmodel includes the brain, skull, meninges, CSF, and superior sagittal sinus (SSS) (**Fig. 2**). The brain is divided into primary structures of cerebral gray matter (GM) (i.e., cerebral cortex), cerebral white matter (WM), corpus callosum (CC), brain stem (BS), cerebellum GM and WM, thalamus, and hippocampus. The cerebrum is further divided into frontal, frontal, parietal, temporal, and occipital lobes; CSF is divided into outer CSF and ventricular system including lateral ventricles, 3^rd^ and 4^th^ ventricles connected by the cerebral aqueduct. Continuous mesh is used between brain components throughout the model. The total number of elements in the head model is 4.4 million hexahedral and 0.54 million quad elements. The minimum Jacobian in the brain is 0.45. The brain is modeled as hyper-viscoelastic material to account for large deformations of the tissue, with additional linear viscoelastic terms to account for the strain rate dependence. Pia, dura/falx/tentorium are modeled with nonlinear hyperelastic material using simplified rubber/foam based on the average stress-strain experimental data [69, 70]. The model has been validated against experimental data of close to or injury level brain-skull relative motion, brain strain, as well as intracranial pressure. Details of the model development, validation, and the capacity for studying brain responses under impact can be found in a previous study [35].

**Fig. 2.**
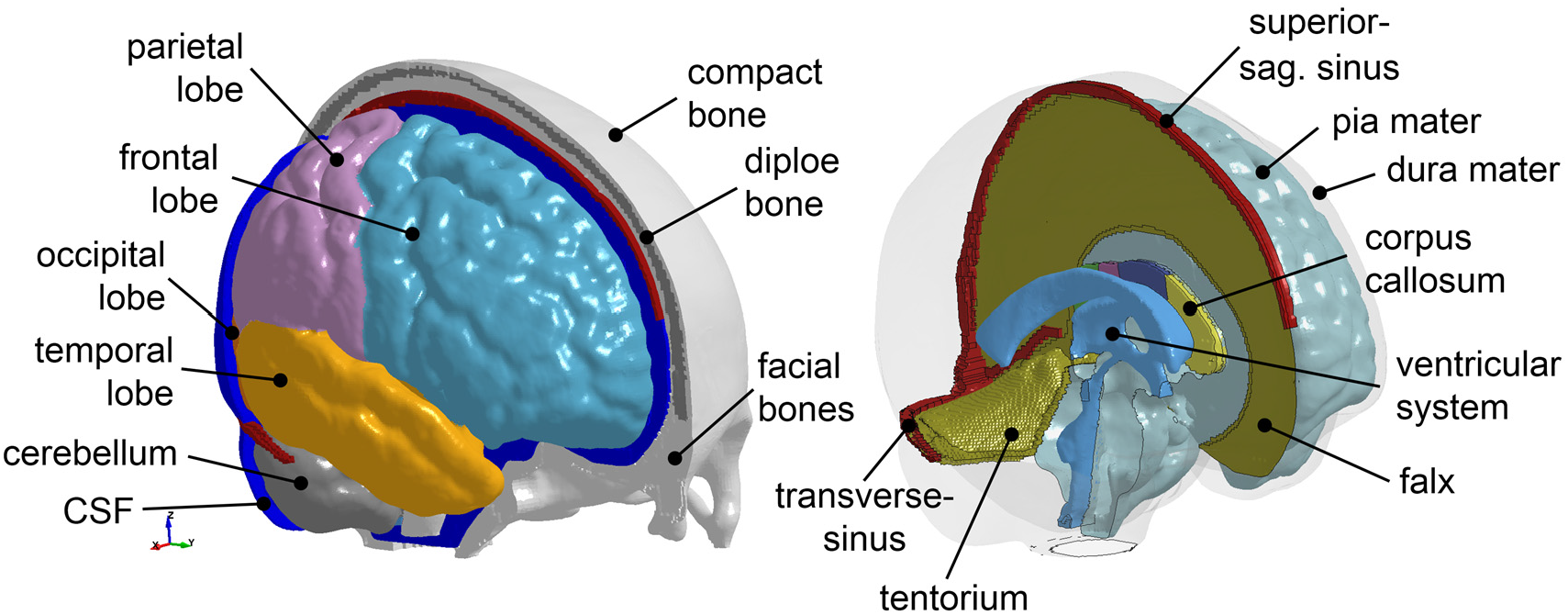
The baseline ADAPT head model with major components illustrated. The meshes are not shown for a better illustration.

### 2.3 Framework for head model personalization from baseline

The framework consists of hierarchical image registrations with multiple features and multimodality images combined with mesh grouping (**Fig. 3**). A complete image registration pipeline is shown at the lower row of **Fig. 3**, with the subject’s image as *moving* image and baseline ICBM as *fixed* image, from which dense displacement fields are obtained (*g*_*demo*_, *g*_*f*1_, *g*_*f*2_…, *g*_*dram*_, *g*_*m*1_, *g*_*m*2_… *g*_*fn*_…). The displacement fields from all steps are added up as *g*_*sub j*_ to morph the baseline ADAPT head model. Afterward, the WM of the morphed brain is regrouped according to the segmented WM image mask from the subject’s T1W image, resulting in the final subject-specific model. The baseline image is warped via inverse of displacement fields from each step, which is then compared with subjects’ image to quantify registration accuracy by DICE, Jaccard index, and HD95. Further details regarding registration algorithms for each step are presented in **Sec. 2.2.1**, followed by morphing and mesh grouping.

**Fig. 3.**
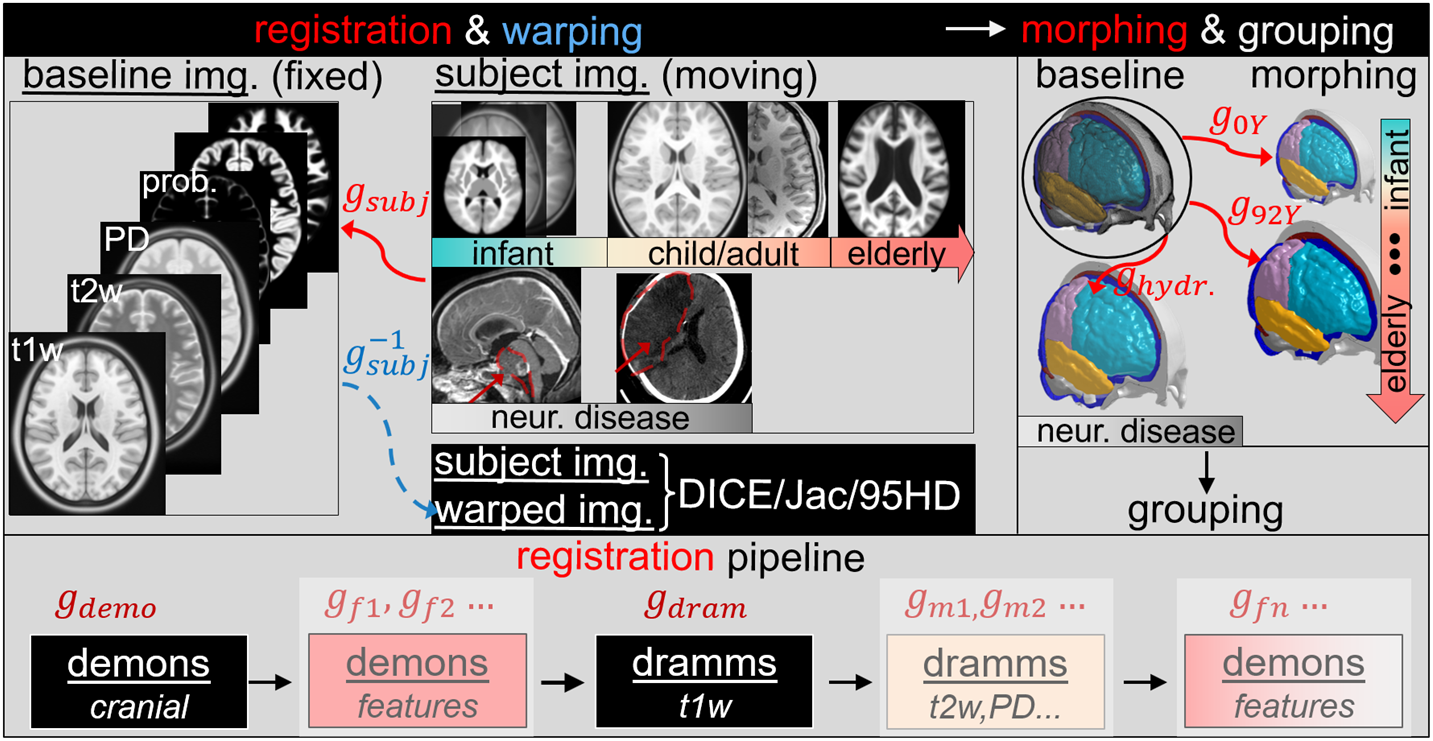
Overview of the personalization framework capable of generating subject-specific head models of infant, child, adult, elderly, and pathological brains with substantial anatomical changes (illustrated by hydrocephalus brain and a brain with decompressive craniotomy). Note that the lower row shows a complete registration pipeline which allows morphing subjects with substantial anatomical changes compared with the baseline ICBM, e.g., a hydrocephalus brain. However, fewer registration steps are sufficient for subjects with small anatomical differences, as shown in Sec 2.5.

#### 2.2.1 Registration with multiple features and multimodality imaging

A complete registration pipeline starts from Demons registration [71] of the segmented cranial volume resulting in a transformation, i.e., dense displacement field *g*_*demo*_, followed by Demons registration of other features (*features* in this study is referred to as the segmented binary image of anatomical regions such as lateral ventricles (LV), corpus callosum (CC) or lesion) obtaining *g*_*f*1_, *g*_*f*2_ etc. Note the input to the feature registration step is a segmented binary image including both the feature and the cranial colume. Thus, voxel-wise registration of the entire brain is performed in this step, and the resulting displacement field captures the local anatomical changes between the baseline and subject image. Afterwards, Dramms registration [72] is then performed with T1W with warped images from Demons steps as input, obtaining *g*_*dram*_. Next, multimodality images, e.g., T2W is then used as input for another Dramms registration for further brain anatomical alignment. Finally, brain lesions can be handled by feature steps using Demons registration.

In particular, for all Demons steps, diffeomorphic Demons registration [71] implemented in the open-source software *Slicer 3D* is used between the segmented cranial masks of the baseline (corresponding to the baseline FE mesh) and the subject after being rigidly aligned using a six-degree-of-freedom rigid registration available in *Slicer* 3D. Dramms registration algorithm [72] implemented as open-source code by the authors (Dramms version 1.5.1, 2018) is performed on the skull stripped MRI images of different modalities inherited from the Demons steps.

The dense displacement fields from all registration steps add up as *g*_*sub j*_ (**Eqn.1**), which are used to morph the baseline mesh for a subject-specific head model as detailed below.

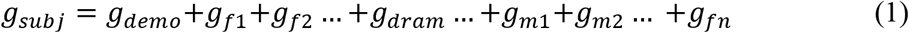

The inverse of displacement fields from each step 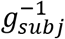 is applied to warp the baseline image *img*_*baseline*_ (**Eqn.3**) (corresponds to the baseline head model), resulting in a warped image *img*_*warped*_ (corresponds to the subject-specific model by mesh morphing).

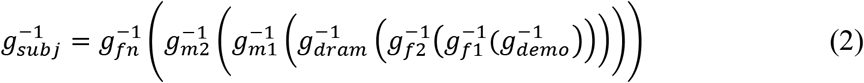

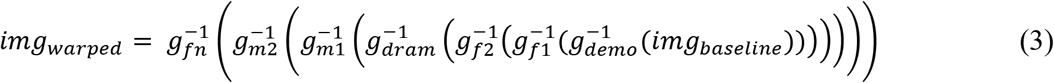

Note that Demons registration allows morphing brains with large differences; however, it tends to result in displacement fields that may lead to excessive element distortion according to our previous experience [73–75]. Therefore, the hierarchical design of the current pipeline is to utilize Demons’ capacity for handling large shape differences by performing Demons registration as the 1^st^ step (*g*_*demo*_) with segmented binary cranial masks as input that allows obtaining a displacement field reflecting well overall cranial shape. Then the 2^nd^ step and last step Demons registration (*g*_*f*1_, *g*_*f*2_… and *g*_*f*n_…) allows capturing large anatomical changes. Dramms registration is performed in the 3^rd^ on the skull stripped T1W images, and 4^th^ step on multimodality images inherited from previous steps. The focus of the Dramms registration steps is to align local brain structures and CSF by using brain MRI information without the need for segmentation. The choice of Dramms is based on its promising performance due to its advancing in hierarchical attribute matching mutual-saliency mechanism [72].

#### 2.2.2 Morphing

As the baseline FE head model (the ADAPT model) is in the same space as the ICBM template image, the resultant displacement fields from all image registration steps capturing the anatomical difference between the subject and the baseline ICBM image are then applied to the baseline head model (**Fig. 2**) to obtain a personalized head model of the subject. For this, the following step is performed to morph the nodes of the baseline head model to new positions according to

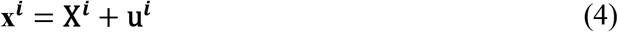

where **X**^*i*^ is the nodal coordinate of node *i*, **u**^*i*^ the linearly interpolated displacement vector at node *n* from *g*_*subj*_, **x**^*i*^ is the updated nodal coordinate, together with the same element definitions as the baseline, forming the subject-specific head model.

#### 2.2.3 Grouping of WM

In order to generate WM subject-specific head model, the morphed brain elements are regrouped based on the segmented binary image of the subject’s cerebral WM obtained using FreeSurfer (see Sec.2.3). Briefly, this step is to assign brain FE elements to WM based on Cartesian coordinates of the segmented WM voxels. The detailed procedures are:

– For each FE element, all WM voxels inside or intersect to a single FE element of the brain are identified based on spatial coordinates.
– The eight vertices and one centroid of each voxel (*i*, *j*, *k*) are judged; a vertice gains a weight of 1 if falling inside the element; the centroid gains a weight of 2 if falling inside the element. Weights of the eight vertices and the centroid of the voxel add up, resulting in a total weighting factor for each voxel *W*_*i*,*j*,*k*_

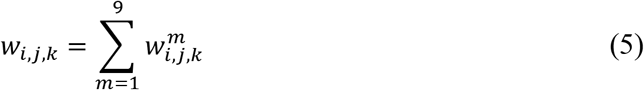
– Finally, weights of each voxel belong to the same label (e.g., the segmented binary image with label A) added up, obtaining a final weight factor for each label. The element is grouped to the label with the largest weight.

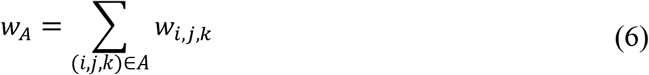

Below shows the regrouped brain elements of the morphed brain using the segmented WM as image label (**Fig. 4**), showing the regrouped WM enclosed by the reconstructed surface of the segmented image of the WM.

**Fig. 4.**
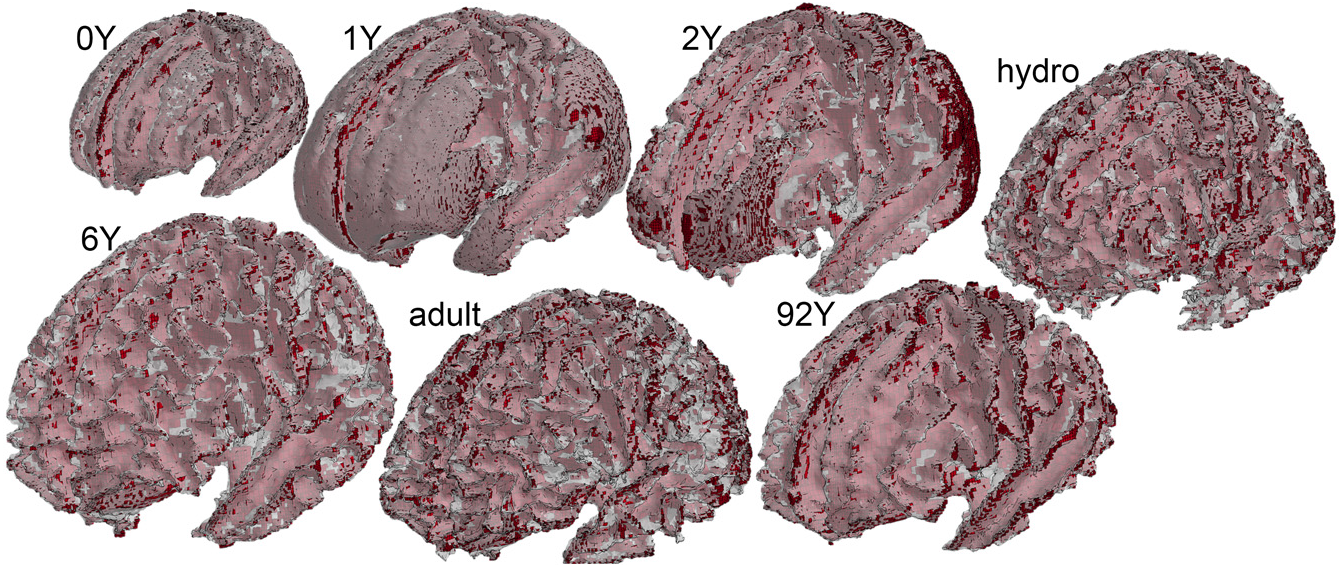
WM regrouped based on subjects’ WM image mask for all the seven subjects. The red color shows the WM elements, and the white transparent shows the surfaces reconstructed from subject’s segmented WM image mask.

### 2.4 Segmentation and evaluation of personalization accuracy

DICE, Jaccard index, and HD95 are calculated as metrics to evaluate registration performance, i.e., how well the personalized T1W image (corresponding to the subject-specific mesh) reflects subjects’ T1W as the ground truth. As the warped image *img*_*warped*_ is directly corresponding to the subject-specific mesh (morphed with the same displacement field). Therefore, these metrics also reflect the personalization accuracy of the generated subject-specific models.

To calculate the matrices, automated segmentation is performed using the software FreeSurfer (version 7.1.0) with the default brain segmentation pipeline (*recon-all*) for both the warped images and subjects’ T1W images without additional manual editing for brain regions. FreeSurfer automatically segmented whole brain and local regions of cerebral GM, WM, CC, BS, hippocampus, thalamus, and cerebellum are used for matrices calculation; while for cranial mask and CC, DICE values are calculated based on semi-automatic segmentation by thresholding followed by noise removal, instead of using FreeSurfer segmentation due to the insufficient quality of segmented cranial mask and CC using *recon-all*. Further, one sagittal slice of CC is used to calculate matrices. Note these segmented labels are only used during DICE calculation, and the quality of the automatic segmentation has no influence on the subject-specific mesh development process.

#### 2.4.1 DICE

DICE is a single metric to measure the spatial overlap between images (Bennett and Miller 2010; Zou et al. 2004), which is defined as twice the number of elements common to both sets divided by the sum of the number of elements in each set:

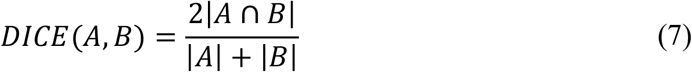

where *A* and *B* denote the binary segmentation labels, |*A*| and |*B*| are the number of voxels in each set, and |*A* ⋂ *B*| is the number of shared voxels by *A* and *B*. The DICE value of 0 implies no overlap between both, whereas a DICE coefficient of 1 indicates perfect overlap.

#### 2.4.2 Jaccard index

Jaccard similarity coefficient defined as the size of the intersection divided by the size of the union of the data sets:

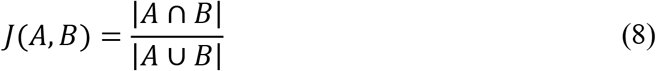

Note Jaccard index is related to DICE, according to *J* = *DICE*/(2 − *DICE*).

#### 2.4.3 HD95

Hausdorff distance (HD95) is defined as

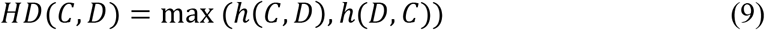

where *C*, *D* are the two sets of vertices from two segmented images

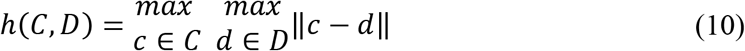

The 95^th^ percentile Hausdorff distance (HD95) is used.

### 2.5 Applications

To demonstrate the capacity of the proposed framework, seven subject-specific head models are generated. As the brain anatomy varies differently compare with the baseline, it is shown, for brains with less anatomical differences compared with the baseline; fewer steps are sufficient. The following shows three typical subtypes of the pipeline with fewer steps (Type I, II, III) as detailed below.

<u>Type I</u> This is basic pipeline contains two steps: Demons registration of the cranial mask and Dramms of the skull stripped T1W image. For the first step Demons registration, the baseline T1W image (i.e., the T1W image corresponding to the baseline FE mesh) and the subject’s T1W are segmented to obtain the cranial mask as input for registration. This two-step pipeline has been shown to achieve good registration accuracy in six adult subjects [35]. The capacity of this pipeline is further demonstrated with a 2Y brain below.

<u>Type II</u> Multiple feature steps are added to the Type I pipeline, allowing align brains with substantial anatomical changes. Below is an application of this pipeline on a hydrocephalus brain, with three feature steps, including LV, CC, and brain lesion. In principle, more feature steps can be added to account for local anatomical changes if needed. Note the lesions in this study refers to mass effect at cranial outers, such as the mass lesion at brain stem front in the hydrocephalus patient, or swelling portion of the brain in patients underwent decompressive craniotomy.

<u>Type III</u> Multi-modality imaging (e.g., T2W image) registration steps are added to Type I pipeline to allow further alignment of brain anatomy. Especially T2W, with higher contrast for CSF/LVs, allows better alignment for brain and CSF/LV.

#### 2.5.1 Infant brain model of 2YO via pipeline Type I: two steps

First, the baseline T1W image (i.e., the T1W image corresponding to the baseline FE mesh) and the subject’s T1W are segmented to obtain the cranial mask, which is used as input for Demons registration from which a dense displacement field *g*_*demo*_ defined at each voxel is obtained (**Fig. 5**). 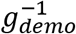 is then used to warp to the baseline T1W image, which is then skull stripped and afterward together with the subject’s skull stripped T1W as input to Dramms registration, obtaining *g*_*dram*_. Finally, the two displacement fields add up, obtaining *g*_*subj*_2*YO*_ = *g*_*demo*_ + *g*_*dram*_. The warped image is obtained via 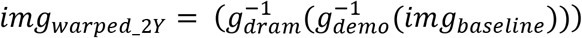, which is compared with the T1W image of the 2Y to evaluate personalization accuracy.

**Fig. 5.**
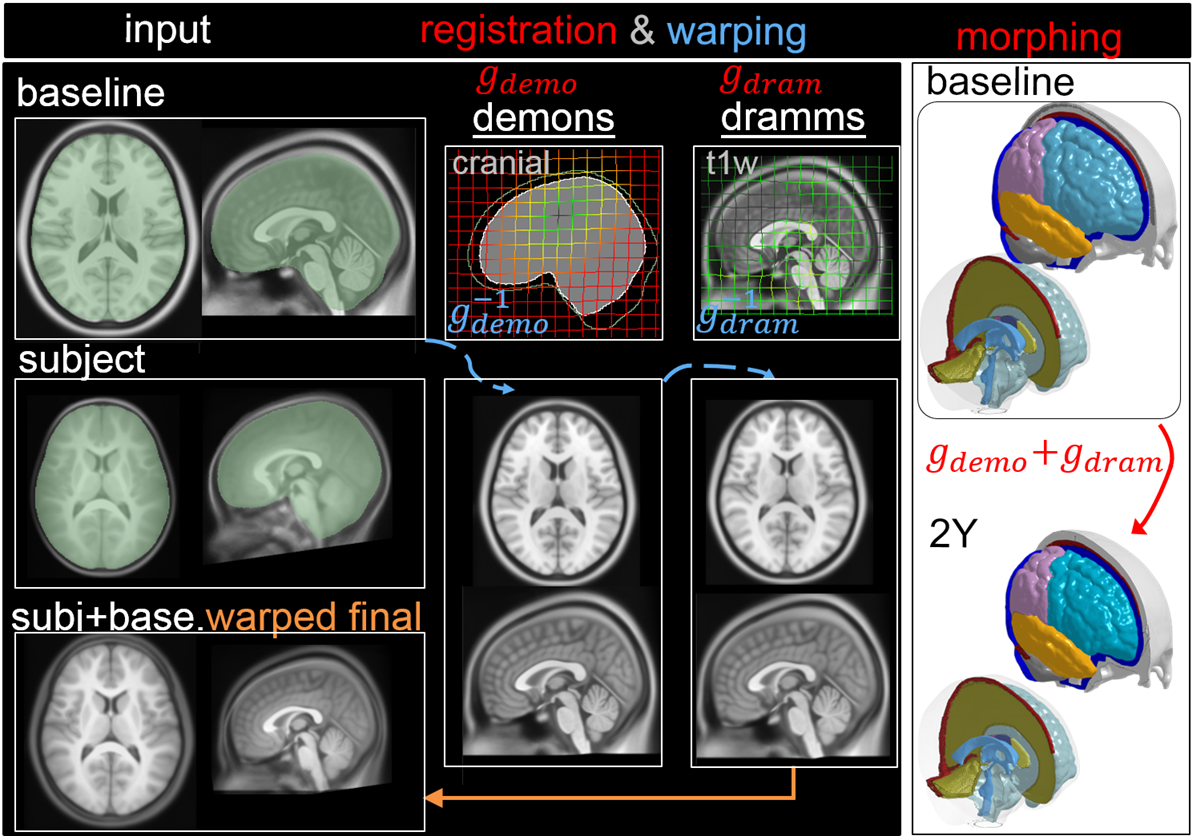
Type I pipeline applied for personalizing the baseline to a subject-specific model of a 2Y, consisting of two steps: (i) Demons registration with cranial mask (ii) Dramms registration with skull-stripped T1W image. The displacement fields obtained via the Demons and Dramms steps *g*_*demo*_ and *g*_*dram*_ are visualized on the grid separately.

**Fig. 6.**
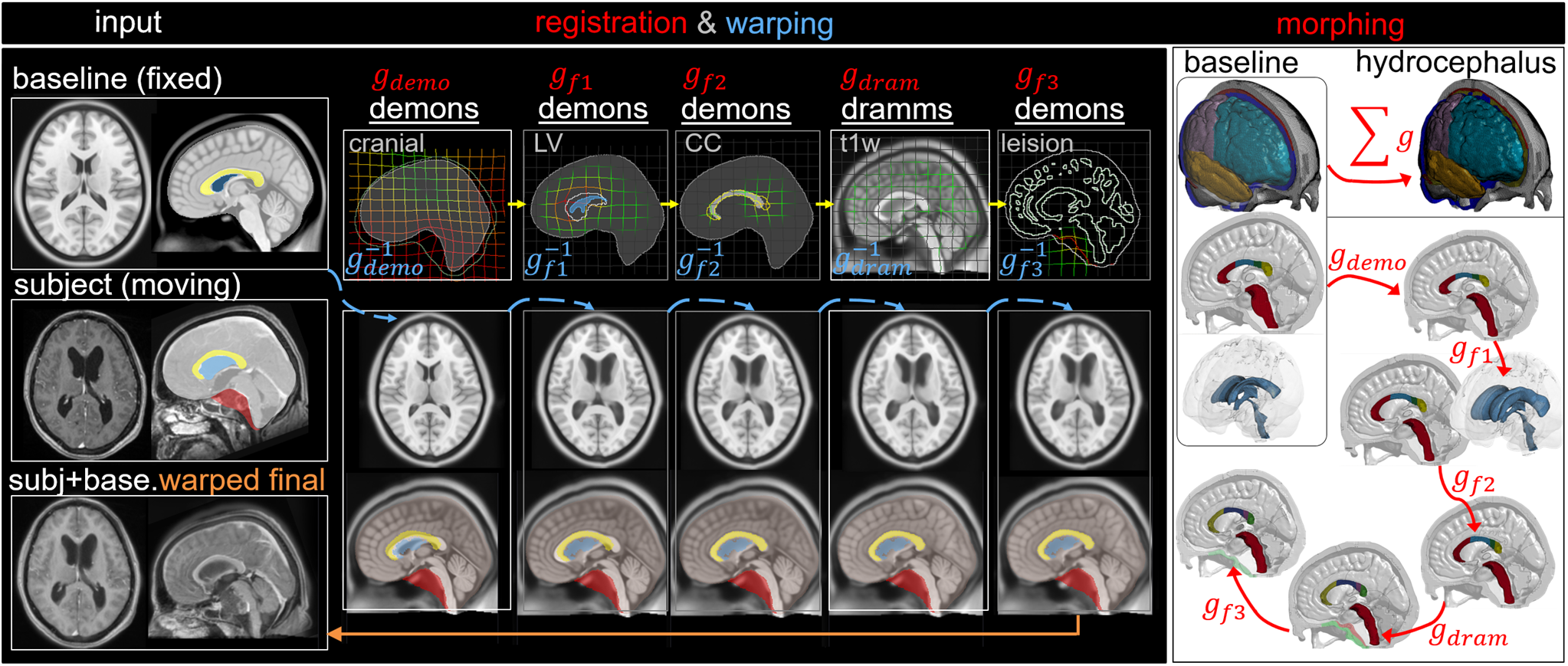
Type II pipeline applied for personalizing the baseline to a subject-specific model of a hydrocephalus brain, consisting of five steps including (i) Demons registration with the segmented cranial mask for capturing the cranial shape and obtain *g*_*demo*_ (ii) Demons registration with segmented LV mask for capturing the enlarged LV and obtain *g*_*f*1_ (iii) Demons registration with segmented CC mask for capturing the CC shape and obtain *g*_*f*2_ (iv) Dramms registration with T1W image for capturing local brain anatomy and obtain *g*_*dram*_ (v) Demons registration with to drag back the skull mesh which is pushed due to the lesion in the cranial mask in step 1 and obtain *g*_*f*3_. The resultant displacement field at each step is visualized on the grid.

#### 2.5.2 Hydrocephalus model via pipeline Type II: multiple features

The workflow is similar to above just adds three extra feature steps to allow capturing the enlarged LV, deformed CC, and brain lesion, resulting in five dense displacement fields add up *g*_*subj_hydro*_ = *g*_*demo*_+*g*_*f*1_+*g*_*f*2_+*g*_*dram*_+*g*_*f*3_. The warped image is obtained via 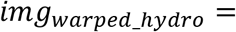 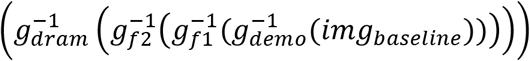, which is in the same space as the subject-specific head model. The final warped image is compared with subject’s T1W to evaluate registration accuracy.

#### 2.5.3 Adult brain model via pipeline Type III: Multimodality

The workflow is similar to Type I just adds one multimodality step of T2W to further align the lateral LVs resulting in three dense displacement fields added up as *g*_*subj_adult*_ = *g*_*demo*_+*g*_*dram*_+*g*_*m*1_, and 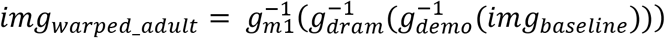 is then used to morph the baseline mesh obtaining the subject-specific head model. The same approach is used to obtain the final warped image by 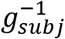 which is compared with the subject T1W to evaluate personalization accuracy.

#### 2.5.4 Pipeline for other subjects including the newborn, 0Y, 1Y, and the 92Y

The 92Y uses Type II pipeline, similar to the hydrocephalus subject, except only one feature step for LV is used, no need for the CC and the lesion feature step as for the hydrocephalus subject, i.e., *g*_*subj_92Y*_ = *g*_*demo*_+*g*_*f*1_+*g*_*dram*_. Interestingly, the Dramms registration captures well the thinning of CC without the CC feature step as for the hydrocephalus subject. It could be due to that the CC of the 92Y, despite showing CC thinning and move upwards due to LV enlargement, is still within healthy development. But the CC in the hydrocephalus patient is arched upwards due to pathologically enlarged LV. This might be due to that the Dramms algorithm captures well normal development of aging, but also could be due to the image contrast of the 92Y comparing with the baseline than the hydrocephalus image.

The 6Y uses only one step Dramms registration since the baseline and the 6Y both aligned to the ICBM template, with very similar brain shape already. Thus, one step Dramms registration achieves good brain anatomy alignment.

The 0Y and the 1Y in principle, could use the same pipeline as the 2Y. However, in this study, an alternative approach is used, by using the 2Y as an intermediate step, i.e., align the 1Y (as *moving* image) to the 2Y (as *fixed* image), obtaining a displacement field *g*_*lY_to_2Y*_, then the final displacement field is used for personalizing the baseline model to the 1Y writes: *g*_*subj*_1*Y*_ = *g*_*subj*_2*YO*_ + *g*_1*Y*_*to*_2*Y*_, and the warped image is obtained via 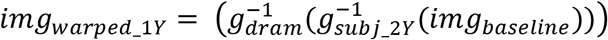 Note the *g*_*sub j*_2*YO*_ contains two parts as described above.

## 3 Results

### 3.1 Personalized head models and element quality

The generated subject-specific head models generated are shown in **Fig. 8**. The pipeline is capable of generating subject-specific head models of widely varying intracranial volume and local brain regions with substantial anatomical changes (**Fig. 9**). Especially, the hydrocephalus has enlarged LV with an Ivan index of 0.36 is captured, attributing the feature step. The element quality for the models is listed in **Table 2**. In the seven subject-specific head models generated, most brain elements (95.9% ± 1.5% on average for the seven subjects) have a Jacobian over 0.5. The minimum Jacobian in the seven head models is all above 0.13 (occurs in the hydrocephalus brain). In this study, the mesh quality is considered to be satisfactory when at least 95% of the elements have a Jacobian over 0.5.

**Table 2.** Element quality of the ADAPT model and the subject-specific head models generated by morphing.

| Head model | Element quality Index |  |  |  |  |  |  |  |  |  |  |  |
| --- | --- | --- | --- | --- | --- | --- | --- | --- | --- | --- | --- | --- |
|  | Jacobian<br>≥0.5 |  | Warpage (°)<br>≤30 |  | Skew (°)<br>≤60 |  | Aspect ratio<br>≤8 |  | Min. angle (°)<br>≥30 |  | Max angle (°)<br>≤150 |  |
|  | percent | min | percent | max | percent | max | percent | max | percent | min | percent | max |
| ADAPT | 98% | 0.45 | 92% | 111.76 | 99.9% | 69.95 | 99.9% | 6.62 | 99.8% | 17.98 | 99.9% | 161.94 |
| Personalized models |  |  |  |  |  |  |  |  |  |  |  |  |
| 0Y | 96% | 0.34 | 92% | 112.60 | 99.9% | 70.93 | 99.9% | 7.99 | 99.9% | 14.58 | 98.0% | 168.41 |
| 1Y | 97% | 0.31 | 92% | 111.99 | 99.9% | 69.68 | 99.9% | 8.20 | 99.9% | 16.23 | 99.0% | 167.85 |
| 2Y | 97% | 0.36 | 92% | 110.26 | 99.9% | 67.19 | 99.9% | 6.64 | 99.9% | 16.39 | 99.0% | 166.38 |
| 6Y | 97% | 0.33 | 95% | 113.52 | 99.9% | 73.38 | 99.9% | 7.61 | 99.9% | 15.65 | 99.0% | 167.18 |
| Adult | 93% | 0.17 | 90% | 139.66 | 99.9% | 74.43 | 99.9% | 10.09 | 99.0% | 8.95 | 98% | 215.91 |
| 92Y | 96% | 0.15 | 94% | 121.81 | 99.9% | 78.53 | 99.9% | 16.09 | 99.0% | 5.56 | 98% | 177.29 |
| Hydrocephalus | 95% | 0.13 | 91% | 118.31 | 99.9% | 77.71 | 99.9% | 11.07 | 99.0% | 7.80 | 97% | 176.79 |

**Fig. 7.**
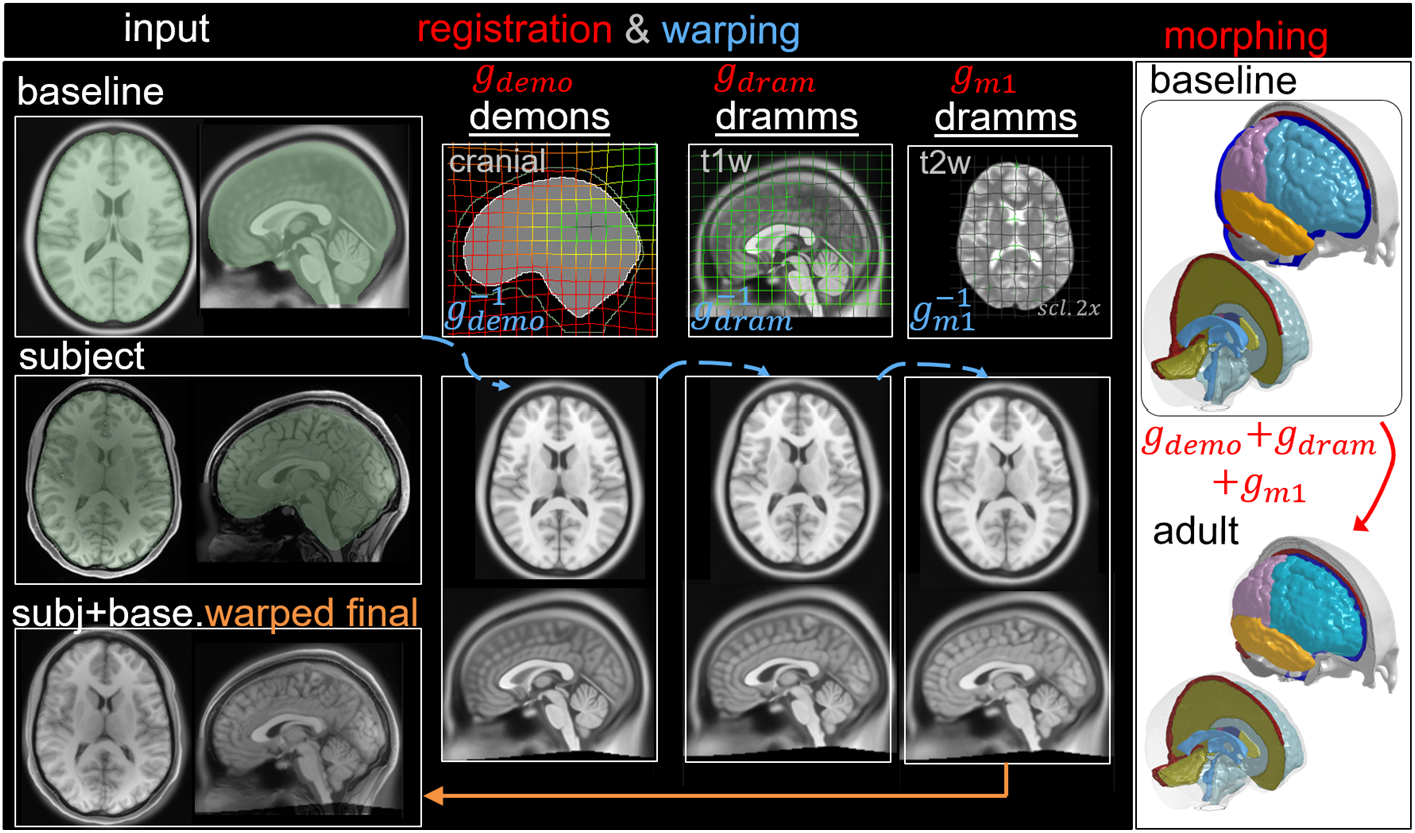
Type III pipeline applied for personalizing the baseline to a subject-specific model of an adult subject, consisting of three steps including: (i) Demons registration with the segmented cranial mask for capturing the cranial shape and obtain *g*_*demo*_ (ii) Dramms registration with T1W image for capturing local brain anatomy and obtain *g*_*dram*_ (iii) Dramms registration with T2W image for capturing align further brain anatomy and LVs and obtain *g*_*m*1_. The resultant displacement field at each step is visualized on the grid.

**Fig. 8.**
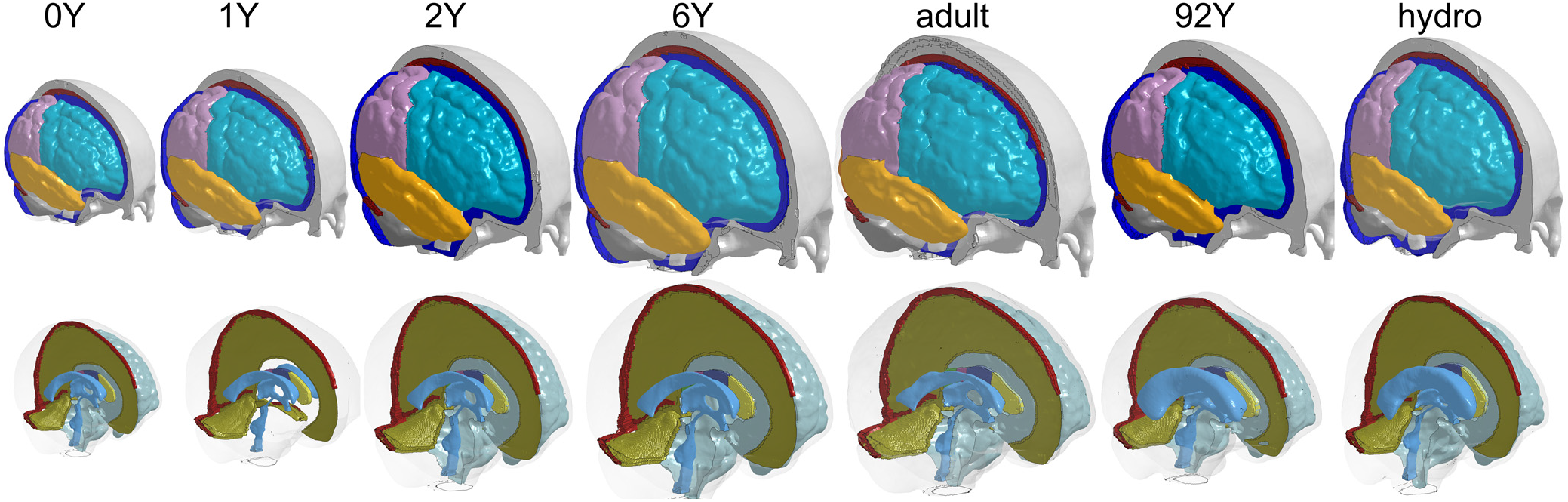
Seven subject-specific head models generated including 0Y, 1YO, 2YO, 6YO, adult, 92YO, and a hydrocephalus brain (the same length scale applies).

**Fig. 9.**
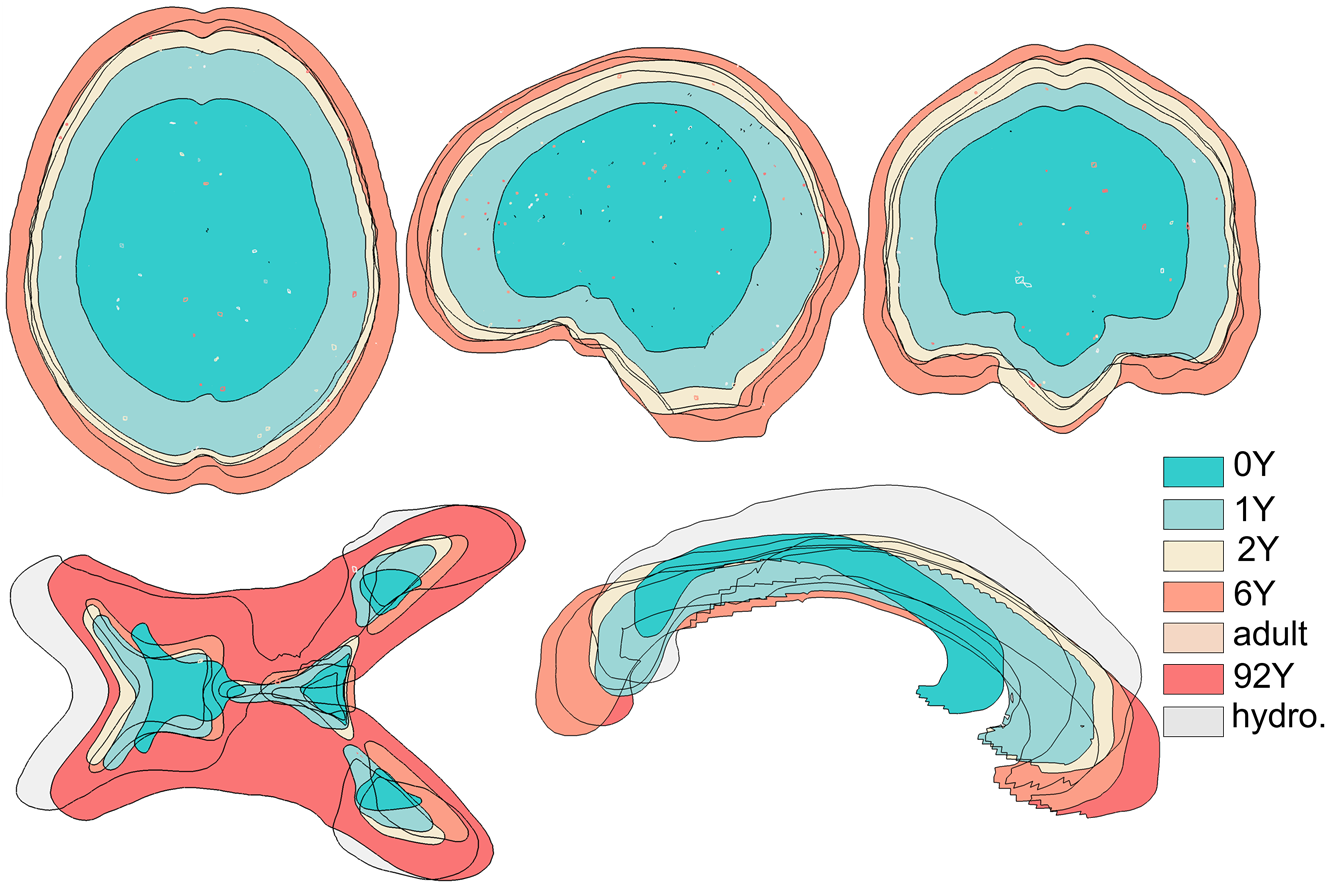
Seven models aligned together, showing the capacity of the framework allowing generating personalized head models with widely varying intracranial volume (upper row) and local brain structures of lateral ventricles and corpus callosum (lower row).

### 3.2 Personalization accuracy

The baseline ICBM image (**Fig. 10**, a) corresponding to the baseline FE head model is warped to subjects (**Fig. 10**, b) and compare with subjects’ image as golden truth (**Fig. 10**, c). The warped images and subjects’ images are segmented using FreeSurfer for registration accuracy evaluation. Note cerebellum for 0Y subject image has been stripped. Thus, the subsequent evaluation of registration accuracy is without these regions. The segmented binary masks of cranial, brain, and seven local brain regions from the warped T1W images of all brains are overlaid to the golden truth (i.e., the subject’s image), showing the registration accuracy (**Fig. 11**, upper). The boxplot of the DICE, Jaccard index, and HD95 are presented (**Fig. 11**, lower) with exact values provided in tables (**Table 3** - **Table 5**), showing average DICE scores are all >0.9 for the cranial mask, the brain, cerebellum, CC, being 0.97, 0.92, 0.90, 0.94, respectively. The average DICE score for LV is 0.82. The DICE values are comparable to or higher than that in the neuroimaging field [76] despite the higher requirement on the smoothness of displacement field for satisfactory element quality in the personalized head FE models.

**Table 3.**
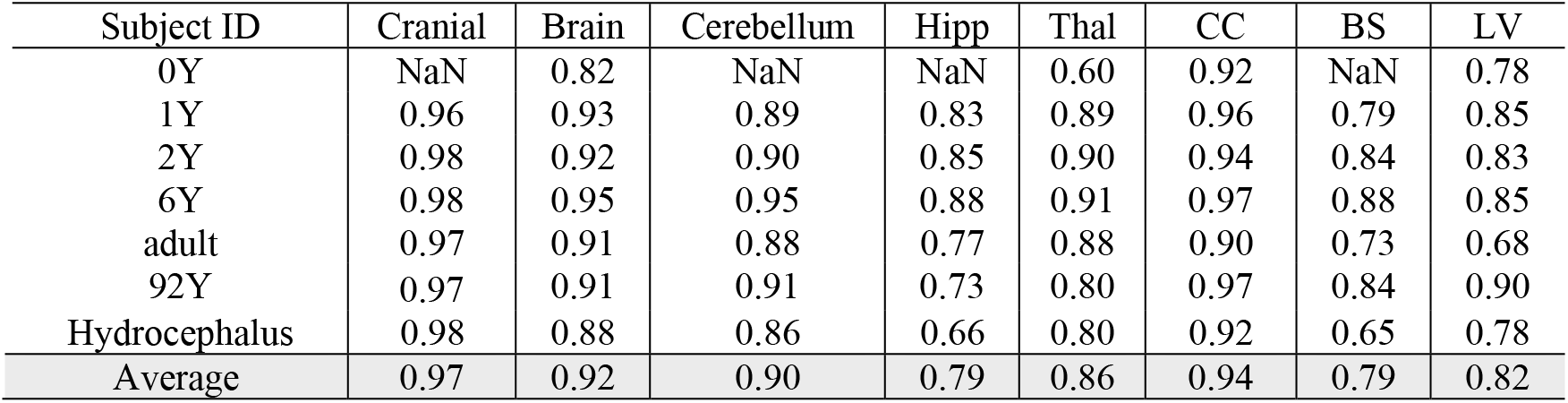
DICE coefficients for the seven subjects.

**Table 4.** Jaccard index for the seven subjects.

| Subject ID | Cranial | Brain | Cerebellum | Hipp | Thal | CC | BS | LV |
| --- | --- | --- | --- | --- | --- | --- | --- | --- |
| 0Y | NaN | 0.70 | NaN | NaN | 0.43 | 0.85 | NaN | 0.63 |
| 1Y | 0.92 | 0.87 | 0.80 | 0.72 | 0.79 | 0.92 | 0.65 | 0.74 |
| 2Y | 0.96 | 0.85 | 0.81 | 0.74 | 0.81 | 0.89 | 0.72 | 0.71 |
| 6Y | 0.97 | 0.90 | 0.91 | 0.79 | 0.83 | 0.95 | 0.79 | 0.74 |
| adult | 0.94 | 0.84 | 0.78 | 0.62 | 0.78 | 0.82 | 0.57 | 0.51 |
| 92Y | 0.95 | 0.83 | 0.84 | 0.57 | 0.67 | 0.94 | 0.72 | 0.82 |
| Hydrocephalus | 0.97 | 0.79 | 0.75 | 0.49 | 0.67 | 0.85 | 0.49 | 0.64 |
| Average | 0.95 | 0.85 | 0.82 | 0.66 | 0.76 | 0.89 | 0.66 | 0.69 |

**Table 5.**
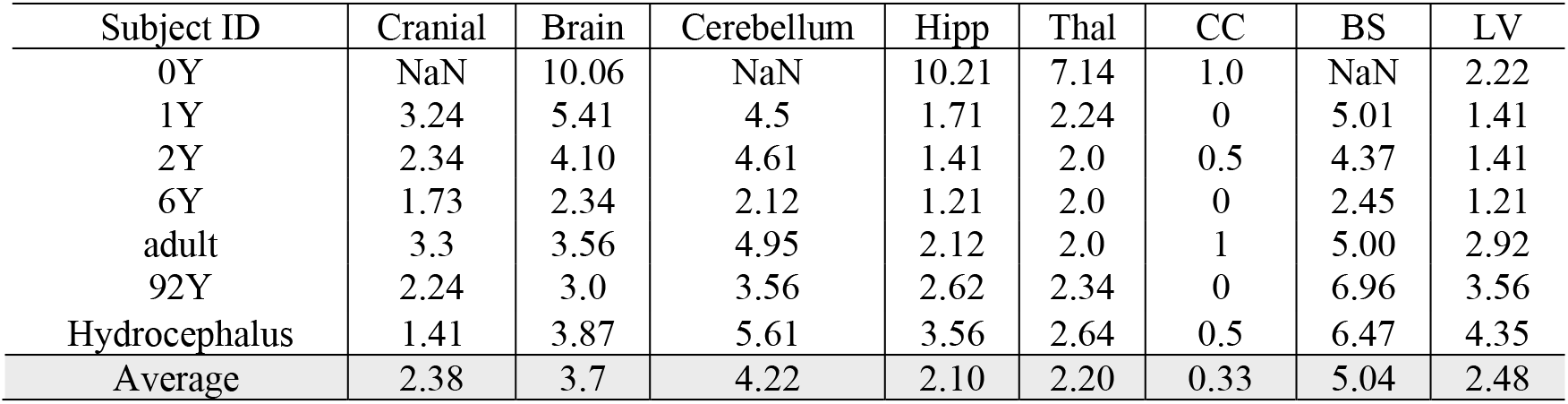
HD95 for the seven subjects.

**Fig. 10.**
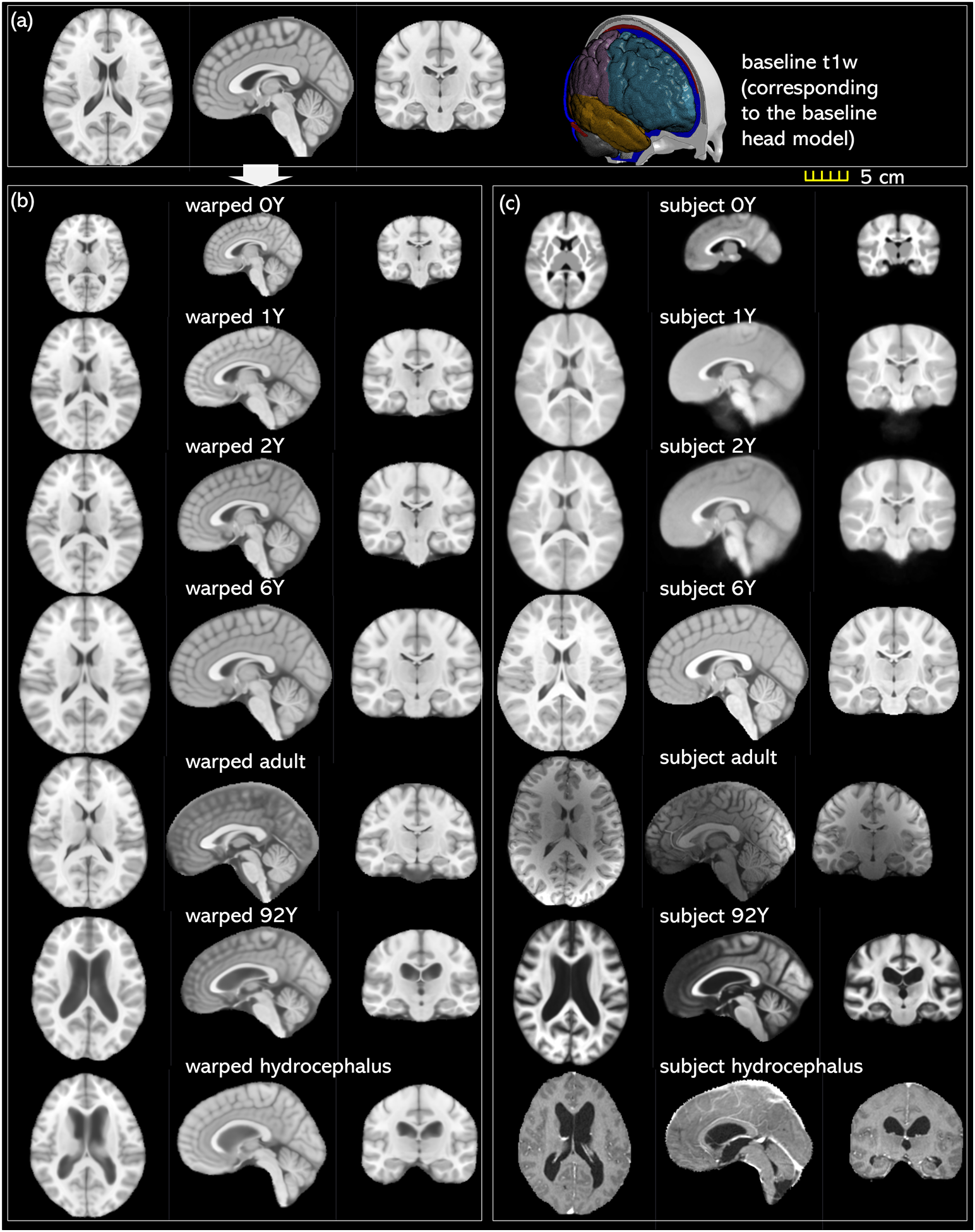
Baseline T1W image of the ICBM template **(a)** and personalized T1W image **(b)** comparing with the ground truth of the T1W image of the subject **(c)**. Transverse, sagittal, and coronal cross-sections are captured for each brain.

**Fig. 11.**
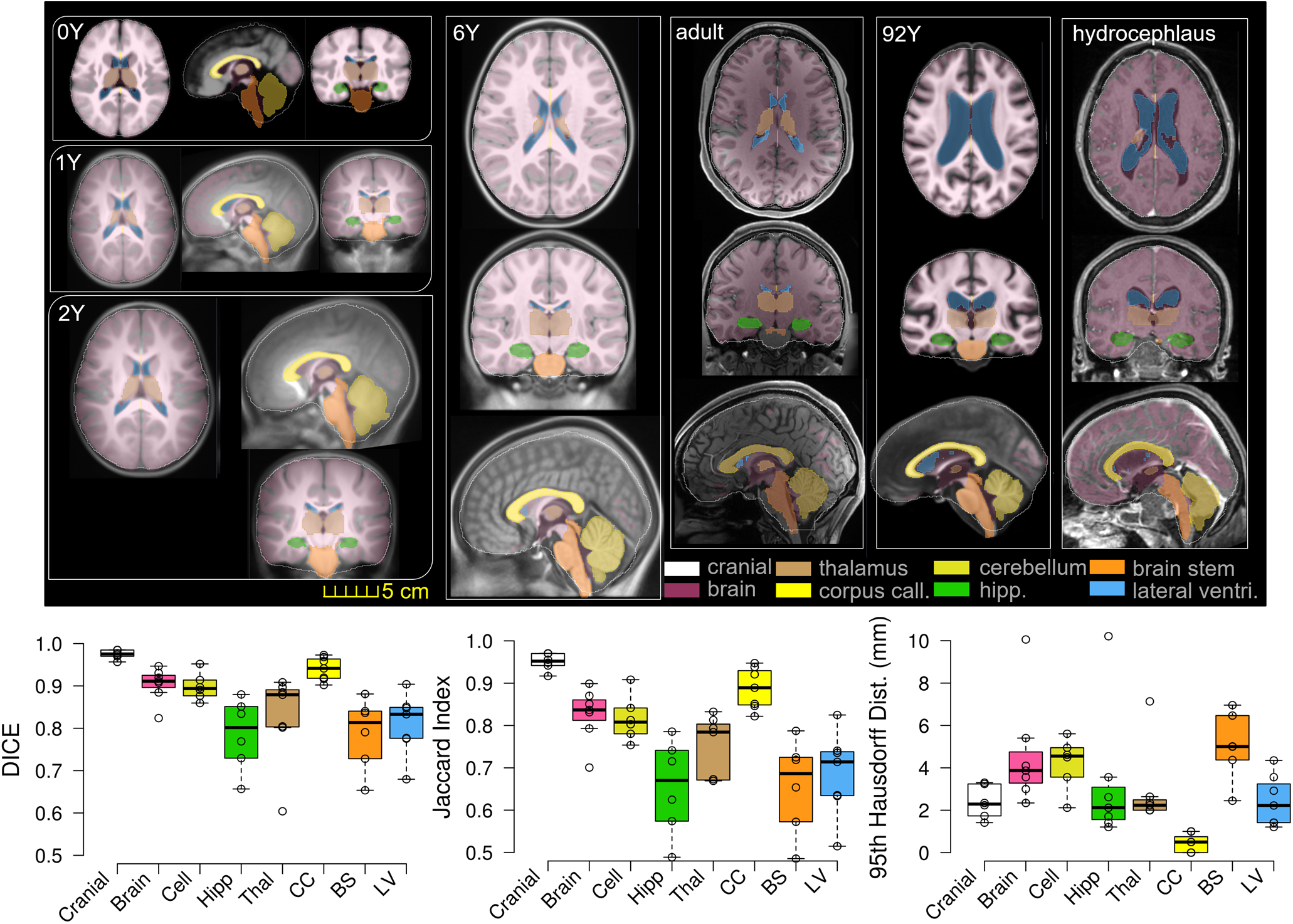
Segmented labels of brain components for matrices calculation for the seven subjects, including the cranial mask, the brain, and local brain regions of cerebrum GM, WM, cerebellum, hippocampus, thalamus, CC, BS, as well as lateral ventricles. T1W image of the subject (golden truth) is overlaid with the segmented regions from the warped T1W image. Boxplot of DICE, Jaccard index, and 95HD. The boxplots show the median, minimum, and maximum values.

### 3.3 Hydrocephalus and the elderly brain: Importance of the feature step and the higher reequipment on displacement smoothness for mesh morphing

The mesh after each morphing step shown in **Fig. 12** illustrates the importance of the feature steps which allow capturing subject’s cranial shape (**Fig. 12**, a), enlarged LVs (**Fig. 12**, b), CC (**Fig. 12**, c), as well as pushing back of the skull mesh (**Fig. 12**, f), while local brain structures are captured by Dramms registration (**Fig. 12**, d). The meshes are morphed from the baseline with displacement fields obtained via the image registration pipeline shown in **Fig. 6**.

**Fig. 12.**
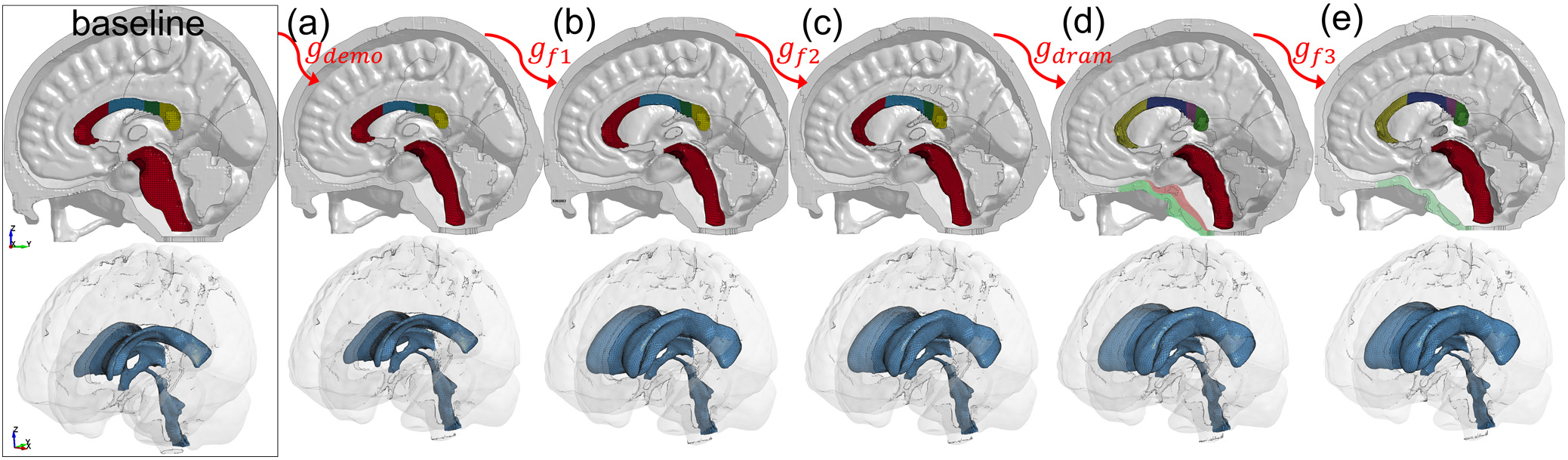
Morphed FE meshes after each step of the five steps for the hydrocephalus subject.

To further illustrate its importance of the feature registrations, a parametric study is performed for the 92Y brain by excluding the LV feature step, i.e., a complete pipeline for the parametric study writes *g*_*subj*_92*Y*_ = *g*_*demo*_+*g*_*dram*_ and 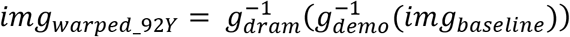. The Results show Dramms registration, even using the largest allowable smoothing factor (g = 1), leads to FE mesh with negative Jacobian in some elements. For example, one FE element at the frontal horn of the LV with a Jacobian of 0.5 in the baseline mesh (**Fig. 13**, right upper, A), when morphed by the parametric pipeline (**Fig. 13**, upper), resulting in a negative Jacobian J = −0.05 (**Fig. 13**, right upper, B). In contrast, the same element, when morphed by the original pipeline with three steps (**Fig. 13**, upper), ended in J = 0.31. Worth noting that pipeline without the LV feature step not only resulting in FE elements with negative Jacobian preventing from simulations being performed, but the registration accuracy is also diminished, as the warped image has a much smaller ventricle (**Fig. 13**, b) comparing achieved by the three-step pipeline (**Fig. 13**, a). This example also demonstrates the higher requirement on the smoothness of displacement than in the neuroimaging field when only Jacobian of the displacement field is of concern, with a more detailed analysis provided in **Appendix 2.**

**Fig. 13.**
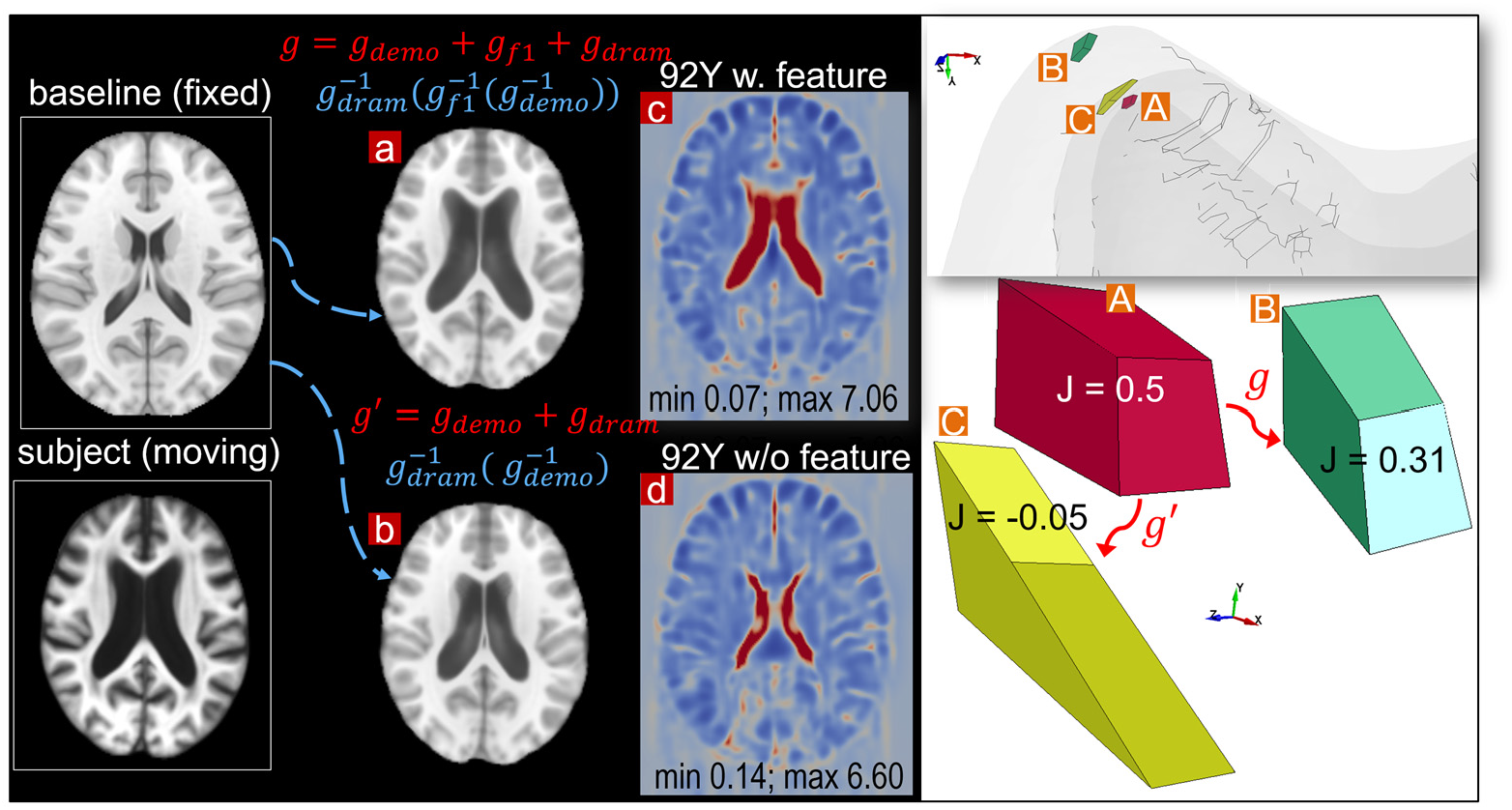
Parametric pipeline for the 92Y without the lateral ventricle feature step **(b)** comparing with the default pipeline **(a)**. The parametric pipeline leads to negative Jacobian in some elements in the personalized mesh (right figure), despite all Jacobian determinant of the obtained displacement field (all steps add up) are all positive. One representative axial slice of the Jacobian determinant is shown with the minimum, and the maximum value in the entire brain indicated **(c, d)**.

## 4 Discussions

This study presents a personalization framework for efficient generation of subject-specific head models, demonstrated by seven brains within the whole lifespan and a hydrocephalus brain with substantial anatomical changes. The approach shows competitive registration accuracy despite a higher requirement on the smoothness of displacement field concerning element quality of the morphed mesh, attributed to the multiple features steps using Demons registration providing a proper initialization for subsequent Dramms registration of T1W and multimodality for further alignment of the brain structures. As a final step of the framework, mesh grouping of brain regions according to subject’s image mask allows subject-specific brain regions to be incorporated accurately, as exemplified with WM, to compensate for the rapid growth in WM/GM in early infancy. The promising performance shows that the framework has the potential to personalize the baseline head model to almost any brains with substantial changes, resulting in valid FE models ready for personalized FE simulations without manual repairing. To the knowledge of the author, this is the first study aligning such a broad scope of brain images suitable for mesh morphing. The personalization framework and the generated models open opportunities for personalized simulations in many fields within neurosciences, especially in the biomechanics of TBIs, in which personalized simulations are hindered due to the meshing challenge. The personalization framework is equally applicable for head models for other applications, such as personalized neurosurgical simulations and brain stimulation.

The efficiency of the hierarchical two-step pipeline combining Demons and Dramms (Type I) has been previously validated in six healthy adult subjects standard T1W MRI with high quality [35]. In this study, an extended framework is proposed capable for more challenging situations for obtaining high-quality alignment across heterogeneous data of whole lifespan and pathological brains with substantial anatomical changes by introducing multiple feature steps as demonstrated with the hydrocephalus (**Fig. 12**) and the 92Y brain (**Fig. 13**). The registration accuracy for the more challenging situations is comparable with the six healthy adults, with average DICE scores for the cerebellum, CC, and brain all above 0.9. Notably, the average DICE score for LV for the seven subjects in this study is 0.82, higher than the average value of 0.71 for the six adult subjects [35]. Note in this study, the same adult subject (Subject ID 771354) is used here by adding an extra T2W MRI registration step (Type III), allowing better alignment of the brain and CSF/LV comparing with previous results without this step (see **Appendix 1**). The framework also contains a final mesh grouping step that allows accounting for subject-specific WM accurately in infants due to growth, not possible for registration to capture, while all other brain regions such as CC with good registration performance not need mesh grouping. Thus, the promising performance supports the idea that the framework may be possible to personalize the baseline model to almost any brains with substantial anatomical changes, besides the hydrocephalus brain as demonstrated, brains with other structural changes such as decompressive craniotomy with brain expanded outside the skull [75] can also be achieved.

Inter-subject registration between brains with substantial anatomical differences is still challenging with limited registration performance [62]. Worth noting that, image registration when applying registration for FE morphing, there is an extra challenge on the smoothness of displacement fields for acceptable element quality of the morphed mesh. Briefly, higher registration accuracy tends to lead to a worse element quality. Thus, it is a tradeoff between registration accuracy and FE quality. Especially, an FE element becomes invalid if with Jacobian negative, and head injury models with negative Jacobian elements are not runnable in most FE analysis software. Even runnable, it will lead to inaccurate and unacceptable computational results due to its ignorance of correct energy. While in the neuroimaging field, for physically plausible morphing, only the Jacobian determinant of displacement field is to be ensured, which is often a looser requirement than FE Jacobian (see detailed analysis in **Appendix 2**). Despite the higher requirement, this study achieves competitive results and higher compared reported in the neuroimaging field. For example, the Jaccard index for inter-subject registration reported in a previous study using popular deformable registration algorithms reported almost all below 0.6 for all brain regions [76]. In contrast, the current study archives average for all regions, all above 0.66. Worth note this study chooses to evaluate DICE for CC on one sagittal slice due to the inaccuracy of CC segmentation from FS, which may make the values for CC not be comparable with others. While the index for other components is comparable as these components are all from FreeSurfer without any manual editing.

The applications of the proposed personalization framework show that different types of pipelines can be used depending on brain anatomy differences between the subject and template image, as well as the subject’s image quality and available imaging modalities. For brains similar to the baseline, even one step is sufficient, e.g., the 6Y; while for the hydrocephalus brain, five steps are needed to achieve a proper alignment; when T2W images are available, multimodality image allows better alignment of brain and CSF/LVs. In principle, more multimodality registrations can be performed if available from the subject as the baseline image contains imaging modalities of T1W, T2W, PD, and tissue probability maps. Besides, more feature steps can be introduced to handle even more challenging cases. Choosing the proper pipeline for a specific case needs trial and error. However, an overall guideline would be to start from Type I then move on to Type III until achieving satisfactory results. Worth noting, the multiple feature steps can be combined to a binary image with multiple labels and perform at once; however, one feature in each step, as done in this study, tends to be more robust. Further, the framework though demonstrated with a baseline head model with conforming hexahedral elements, it is equally applicable for personalizing other head models as a baseline, e.g., models with tetrahedral elements as commonly used for tDCS, TMS, TUS. The framework is also applicable to smoothed-voxel brain models, as the morphing is to apply the obtained displacement fields to a cloud of nodes, justifying its broad applicability.

Comparing with existing studies registering adult brains, there are fewer studies to align infant brains, which are more challenging partially due to the rapid development of brain anatomy within the first years, especially T1W images are inversed with densities. Not only more challenging for registration algorithms, the evaluation of performance is also more difficult as most segmentation algorithms are developed based on adult images, such as FreeSurfer. Worth noting that the lowest registration accuracy in all brains is for the thalamus in the 0Y; a visual check shows FreeSurfer automatically segmented thalamus not accurate enough. Future studies can employ infant Freesurfer [77] for more accurate segmentation for infant brain images, thus allows more objective evaluation of personalization accuracy. For adult brain mesh morphing, a recent study presents an image registration-based approach showing promising for morphing smooth-voxel healthy adult brain FE models, using affine and deformable registration algorithms implemented in Advanced Normalization Tools (ANTs) [61]. However, the personalization accuracy hasn’t been quantified. Further, as the authors discussed that pipeline cannot account for brains with large anatomical changes. Approaches allow generating brain models with large anatomical changes such as for hydrocephalus brain and elderly brain as in this study, provides a vast opportunity for studying age-specific and groupwise TBIs. Especially, brains with neurological diseases such as hydrocephalus brain with exceptionally enlarged LVs mimicking the elderly brain may provide a possible clue for new insights into the TBIs. The approach also opens the opportunity for studying how an already vulnerable brain, e.g., a hydrocephalus patient, may sustain a TBI injury risk under fall impact, especially hydrocephalus patients who are more prone to fall [78]. Until today, the biomechanics of TBIs in these groups are much understudied, partially due to the meshing challenge.

Comparing with many existing studies of TBIs for healthy adults, the injury mechanisms of infant and children are also understudied, being only a handful of child/infant head models today, e.g., [41–43]. In addition to the meshing challenge for adult models, the development of additional unique features of suture and fontanel plays essential roles in head impact response [43]. Previously, mesh morphing has also been used for morphing a baseline infant head model to different ages using Radial basis function (RBF) to interpolate the displacement field obtained from land markers the anatomical features of suture and skull surface [42]. Unlike the image registration-based morphing, the RBF approach needs manual indentation of land markers, which is often tedious [60]. The RBF approach also does not account for brain anatomies. Comparatively, the morphed detailed infant brain models in this study, when combined with the detailed skull and scalp models [43, 45] will allow studying brain injury [78] biomechanics under impact for infant head model for abusive head trauma with important legal applications for forensic diagnosis. The newborn infant head models may be used for studying delivery related neurotrauma and studying new intervention approaches for clinical problems.

Some limitations and future works to be mentioned. First, the proposed framework allows efficient generation of subject-specific head models with competitive personalization accuracy and satisfactory element quality without mesh repairing. However, The steps are a bit tedious, especially for the brains with large anatomical changes with many steps involved, though the steps can be automated. However, comparing with the efforts takes for generating subject-specific head models (weeks or even months), this effort is considered minimal. Second, the current framework captures well brain internal structures, but the major sulci and gyri lines are not evaluated like most studies in the neuroimaging field. Third, Dramms registration algorithm is chosen for registering brain MRI images in this study; other algorithms such as implemented in ANTs [79] and DARTEL [71], when used within the current framework, may achieve similar performance and is yet to be investigated. Finally, the framework can be extended to include more steps and more advanced nonlinear registration algorithms if needed for even more challenging situations.

## 5 Conclusions

This study proposes a framework for head model personalization by combining hierarchical multiple feature and multimodality imaging registrations with mesh grouping. The framework allows efficient generation of subject-specific head models within the whole lifespan and brains with substantial anatomical changes, thus, facilitates personalized simulations within many neuroscience fields, such as biomechanics of TBIs and brain stimulation. The personalization framework allows individual TBIs to be better studied, which has not been possible so far partially due to the meshing challenge for subject-specific model generation. Further, the generated family of lifespan head models, especially the newborn and elderly, are ready to be used for studying age dependent TBIs and clinical applications.

## Acknowledgments

This work benefits greatly from the open-source imaging dataset and registration algorithms within the neuroimaging field. The WUM HCP data of the adult subject used in this study were provided [in part] by the Human Connectome Project, WU-Minn Consortium (Principal Investigators: David Van Essen and Kamil Ugurbil; 1U54MH091657) funded by the 16 NIH Institutes and Centers that support the NIH Blueprint for Neuroscience Research; and by the McDonnell Center for Systems Neuroscience at Washington University. The computations were performed on resources provided by the Swedish National Infrastructure for Computing (SNIC) at Center for High Performance Computing (PDC) at KTH. The present study was supported by research funds from KTH-Royal Institute of Technology, Stockholm, the Swedish Research Council Grants (nr. 2016-04203, and nr. 2020-04724).

## Appendix 1 Adult subject with multimodality registration

Multimodality T2W Dramms registration improves registration accuracy for local brain regions as demonstrated with the adult, especially the DICE score for LV increases from 0.62 to 0.68 (**Fig. A1**).

**Fig. A1.**
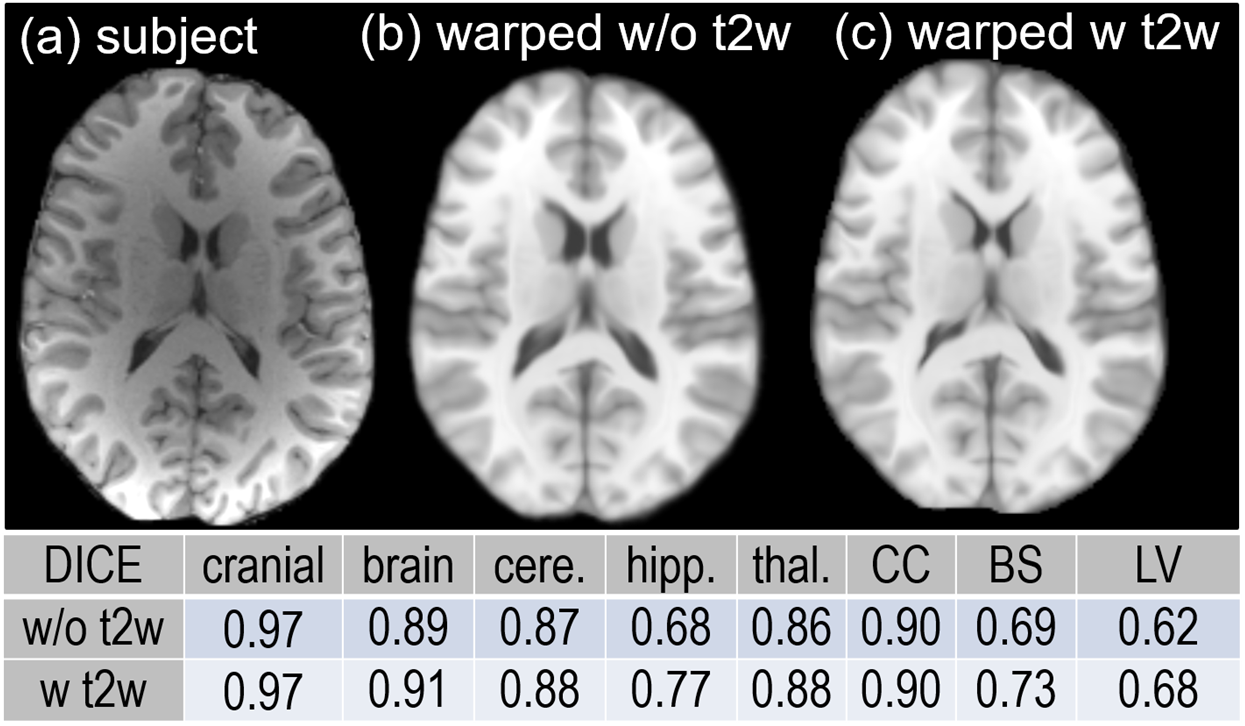
An axial slice of the subject image **(a)** is compared with the warped images without T2W **(b)** and with T2W step **(c)**, showing improved registration accuracy, especially for LV.

## Appendix 2 Higher requirement on displacement smoothness for mesh morphing than neuroimaging field

A higher requirement is posed on the smoothness of the resultant displacement field when applying for mesh morphing, requiring the FE element with positive Jacobian. Note that the terminology Jacobian is used both in FEM also in the neuroimaging field. In FEM, Jacobian is used describe element quality and indicates how an element defined in global coordinate system (**X**) derivates from ideal case defined in the natural coordinate system (**r**), defined as [80]:

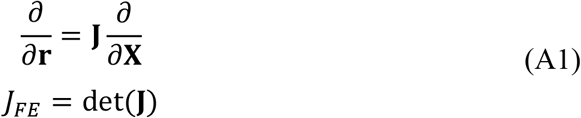

Where **X** and **r** denote the global and natural coordinate system, respectively, **J** is the Jacobian matrix, and *J* is the Jacobian determinant, which is scalar and is often abbreviated as Jacobian. Note that for linear hexahedral elements, *J*_*FE*_ varies inside an element according to the shape function. However, most commercial software reports a single scalar for one element. For example, Hypermesh evaluates the determinant of the Jacobian matrix at each of the element’s Gauss integration points or at the element’s corner nodes and reports the ratio between the smallest and the largest. The Jacobian ratio (also abbreviated as Jacobian) ranges from −1 to 1, and a perfectly cubic hexahedral element has *J*_*FE*_ = 1. The elements with *J*_*FE*_ < 0 represents a concave shape (e.g., element shown in **Fig. 13**) and are prevented from running many FE software.

Jacobian is also used in the neuroimaging field defined as the determinant of the Jacobian matrix of the deformation field as in Eqn. (A2) related to the resultant displacement field from image registration, defined as:

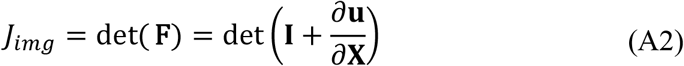

Where **X** is the global coordinate system, **F** is the deformation field, **I** is unit tensor, **u** is the displacement field from image registration. *J*_*img*_ represents the volumetric change at a voxel, with *J*_*img*_ > 0 indicating one-to-one warping between the fix and moving images, which is physically plausible, as shown in **Fig. 13** c, d for the 92Y brain.

The above analysis shows a higher requirement is posed on the smoothness of displacement fields when applying for mesh morphing of FE elements. Since the Jacobian ratio of FE elements (*J*_*FE*_) are usually derivates from the ideal shape and most and *J*_*FE*_ < 1 (values of 0.7 and above are generally acceptable), thus when *u* from registration is imposed to move the element nodes, easier to resulting in elements with negative Jacobian ration, while though the resultant *J*_*img*_ from the same, *u* are all positive, as shown in the example in **Fig. 13**. The Jacobian map of the displacement field added up for mesh morphing for the 2Y, 6Y, 92Y, and the hydrocephalus subjects are presented in **Fig. A2** all calculated on the fixed image, i.e., the ICBM template space.

**Fig. A2.**
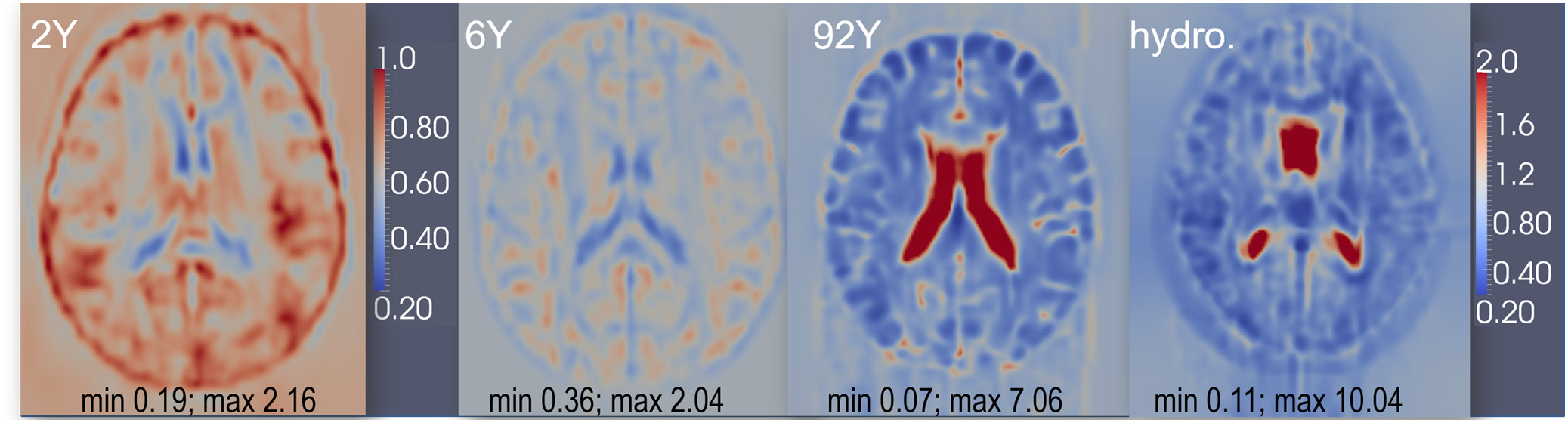
Jacobian determinant of the displacement field obtained from all steps of registration for representative cases of the 1Y, 6Y, 92Y, and the hydrocephalus brain. One representative axial slice is shown with the minimum, and maximum value of the entire brain indicated.

